# H3K4-H3K9 Histone Methylation Patterns and Oncofetal Developmental Networks as Drivers of Cell Fate Decisions in Pediatric High-Grade Gliomas

**DOI:** 10.1101/2024.12.29.630486

**Authors:** Abicumaran Uthamacumaran

**Affiliations:** Department of Experimental Surgery, McGill University, Montreal, Canada; Department of Physics (Alumni), Concordia University, Montreal, Canada; Department of Psychology (Alumni), Concordia University, Montreal, Canada; Oxford Immune Algorithmics, Reading, UK

**Keywords:** Pediatric Gliomas, Phenotypic Plasticity, Cancer Multiomics, Data Science, Systems Medicine, Precision Oncology

## Abstract

This study employs systems medicine approaches, including complex networks and machine learning-driven discovery, to identify key biomarkers governing phenotypic plasticity in pediatric high-grade gliomas (pHGGs), namely, IDHWT glioblastoma and H3K27M diffuse intrinsic pontine glioma (DIPG). By integrating single-cell transcriptomics and histone mass cytometry data, we conceptualize these aggressive tumors as complex adaptive ecosystems driven by hijacked oncofetal developmental programs and pathological attractor dynamics. Our analysis predicts lineage-plasticity markers, including KDM5B (JARID1B), ARID5B, GATA2/6, WNT, TGFβ, NOTCH, CAMK2D, ATF3, DOCK7, FOXO1/3, FOXA2, ASCL4, PRDM9, METTL5/8, RAP1B, CD99, RLIM, TERF1, and LAPTM5, as drivers of cell fate cybernetics. Further, we identified endogenous bioelectric signatures, including GRIK3, GRIN3, SLC5A9, NKAIN4, and KCNJ4/6, as potential reprogramming targets. Additionally, we validate previously discovered plasticity genes such as PDGFRA, EGFR targets, OLIG1/2, FXYD5/6, MTSS1, SEZ6L, MTRN2L1, and SOX11, confirming the robustness of our complex systems approaches. This systems oncology framework offers promising avenues for precision medicine, optimizing patient outcomes by guiding combination therapies informed by single-cell multi-omics and targeting pHGG phenotypic plasticity as therapeutic vulnerabilities. Further, our findings suggest the epigenetic reprogrammability of tumor phenotypic plasticity (i.e., transition therapy) and maladaptive behaviors in pHGG ecosystems toward stable, transdifferentiated states.

## INTRODUCTION

Pediatric high-grade gliomas (pHGGs) represent lethal diseases without any precision diagnostics, effective treatments or prevention (Swanton et al., 2024). These aggressive tumors disrupt developmental processes and tissue homeostasis, leading to aberrant morphogenesis, resistance to therapy, and immune evasion (Senft et al., 2017; Jessa et al., 2019). Central to their pathology is *phenotypic plasticity*—the ability of cells to adaptively transition between lineage identities in response to microenvironmental pressures. This plasticity arises from epigenetic dysregulation, such as oncohistone mutations like H3K27M (H3F3A) and driver mutations like TP53, ACVR1, etc., which destabilize chromatin structure, trapping cells in metastable, multipotent states and impairing their differentiation hierarchy (Shpargel et al., 2014; Paugh et al., 2011; Jessa et al., 2019). In effect, these plastic states foster tumor progression and therapy resistance as *emergent behaviors*, creating an unstable ecosystem of heterogeneous cell populations. Thus, understanding the interplay between differentiation hierarchies, phenotypic transitions, and epigenetic drivers is essential to address the malignant adaptability of pHGGs.

In contrast to their adult counterparts, pHGGs exhibit a heavier epigenetic burden that disrupts normal neurodevelopmental processes (Shpargel et al., 2014; Deshmukh et al., 2022; Sussman et al., 2023; Weller et al.,2024). For instance, diffuse intrinsic pontine gliomas (DIPGs) often originate from oligodendrocyte precursor cells (OPCs) and show transcriptional similarities to the pons and midline structures, while pediatric glioblastomas (GBMs) frequently arise from neural progenitor cells (NPCs) or radial glia (neural stem cells of the SVZ), and exhibit highly vascularized, invasive, and aggressive behavior (Paugh et al., 2011; Deshmukh et al., 2022; Wang et al., 2024). The K27M mutation in DIPG destabilizes chromatin, trapping cells in a developmental arrest, while IDH wild-type (IDHWT) GBMs show fuzzy (hybrid) identities with trilineage cell fate structures steering their epigenetic landscape, and gradients of stem cell markers (Jessa et al., 2019; Couturier et al., 2020; Liu et al., 2022). Emerging evidence suggests that both pHGG subtypes undergo lineage identity transitions between neural stem cells (NSCs; radial glia) and hybrid OPC/NPC states, driven by bifurcations on their developmental (epigenetic) landscape (Wang et al., 2021a,b; Uthamacumaran, 2024). Neuron-glioma interactions and the neuro-metabolic-immune axis further influence these transitions, highlighting the role of epigenetic plasticity as an "evolvability engine" that drives tumor progression (Uthamacumaran, 2024). Despite the cell of origin differences, we hypothesize that similar neurodevelopmental programs (*plasticity networks*) and their state-space attractors (i.e., asymptotic, long-term dynamics) drive the evolution of both glioma subtypes, offering potential targets for ‘transition’ or ‘differentiation’ therapies to constrain their unstable developmental trajectories and guide them towards cell fate commitments (Huang et al., 2005).

Our data science rationale is to address the lack of treatment consensus and unaddressed phenotypic plasticity in pHGG care, focusing on machine learning-driven pattern discovery for precision medicine and paving epigenetic therapies to counter pHGG invasion (Lau et al., 2024). In essence, pHGGs are morphogenetic disorders characterized by microenvironmental evolution, with recent studies using machine learning algorithms and complex network topology to dissect their emergent behaviors (Winkler et al., 2023; Baig et al., 2024; Park et al., 2024). Thus, tumor microenvironment (TME) characterization requires a *whole-systems* perspective, as emphasized by complex systems theory and cybernetics approaches (Winkler et al., 2023; Baig et al., 2024; Park et al., 2024). A complex systems-level approach offers the potential to reveal therapeutic vulnerabilities within the plasticity networks driving these tumors along their neurodevelopmental hierarchy (epigenetic landscape). Thus, *systems medicine* integrates multiomics and machine learning to characterize the *cybernetics* of tumor ecosystems, i.e., the nonlinear feedback loops, interrelationships, and emergent patterns underlying tumor behavior (Uthamacumaran, 2024). By cybernetics, we refer to the control, regulation, and flow of information processing underlying cancer plasticity networks. In our context, cybernetics, derived from the Greek word "*kybernetes*" meaning *to steer*, is the study of how cells navigate or steer their epigenetic landscape through decision-making processes and tumor-microenvironmental networks (Wiener, 1984).

By applying *cybernetics* approaches, we can identify critical regulatory nodes and attractor states steering cell fate choices (Mitchell, 2009; Batterman, 2021). These insights might reveal how tumor cells are developmentally arrested, and trapped in metastable, hybrid identities between NPC and OPC states, halting terminal differentiation (Ocasio et al., 2023). We propose that their phenotypic plasticity arises as an emergent property, reflecting a *maladaptive behavior* or stress response to this differentiation blockade, as they explore various cell fate choices (Jessa et al., 2019; Wang et al., 2024; Uthamacumaran, 2024). Our findings in this study, suggest a gliomagenesis-leukemogenesis developmental axis driven by shared epigenetic and transcriptional networks across pHGG subtypes and pediatric cancer lineages. Specifically, our approaches reveal that cancer stem cells (CSCs) in the pHGG ecosystem show a hybrid spectrum of NSC-derived cell states, including glial stem cells (GSCs), NPCs, OPCs, myeloid, and hematopoietic stem cells (HSC), contributing to invasion, immune evasion, and therapy resistance. Through this transdisciplinary framework, we propose combining data science, dynamical systems approaches, network science, and algorithmic information theory, to identify predictive biomarkers and novel therapeutic strategies that restore neurodevelopmental hierarchies, ultimately mitigating the aggressive behaviors of pHGGs and improving quality patient care (outcomes).

## METHODS

### Gene Expression Counts

Gene expression profiling was performed using Expression Profiling by High-Throughput Sequencing – single-cell RNA sequencing (scRNA-seq), generating count matrices in TPM (transcripts per million). Filtering steps retained cells with a minimum of 200 detected genes and excluded genes expressed in fewer than three cells. The datasets included Neftel et al. (1,943 cells from n=8 (eight) pediatric IDH-WT GBM patients) and Filbin et al. (2,458 cells from n=6 (six) H3K27M DIPG patients) (Filbin et al., 2018; Neftel et al., 2019).

### scEpath

scEpath is an algorithm for reconstructing cellular trajectories and quantifying energy landscapes from scRNA-seq data, inspired by Waddington’s epigenetic landscape (Jin et al., 2018). It computes single-cell energy (scEnergy) using a statistical physics-based model of gene expression and constructs probabilistic transition graphs to map cell state hierarchies and transitions. Leveraging this, we inferred key regulatory transcription factors (TFs) for cell fate differentiation by considering all PDG genes with a standard deviation greater than 0.5 and a Bonferroni-corrected p-value below a significance level of alpha equal to 0.01 for the expression greater than a threshold (e.g., log2 fold-change greater than 1). The probabilistic-directed graph network and cell lineage hierarchy inference parameters were kept at default settings (quick_construct equal to 1; tau equal to 0.4; alpha equal to 0.01; theta one equal to 0.8). Pseudotime-dependent genes were identified using parameters sd_thresh equal to 0.5 and sig_thresh equal to 0.01 (Jin et al., 2018).

### CellRouter

CellRouter is a single-cell trajectory inference tool designed to uncover transition pathways and gene signatures (Lummertz da Rocha et al., 2018). Input gene expression data undergoes preprocessing, including log normalization, scaling, and dimensionality reduction using PCA (50 components, seed=42). Clustering is performed with graph-based methods (k=20, num.pcs=15), followed by t-SNE visualization to define clusters and map population distributions. Subpopulation markers are identified through differential expression analysis with a fold-change threshold of ≥ 0.5. For trajectory inference, KNN graphs (k=10) based on Jaccard similarity are constructed to connect subpopulations, and path-finding algorithms identify dynamic transitions between clusters, ranking trajectories by cost, flow, and path length. Gene regulatory networks (GRNs) are then built using a z-score cutoff of 5, focusing on genes upregulated or downregulated along trajectories. Finally, clusters and pseudotime dynamics are visualized with PCA, t-SNE, and heat maps of trajectory-specific kinetic gene clusters. The SCENIC algorithm was applied only to IDHWT glioblastoma, as the K27M lineage development identity has been well established (Jessa et al., 2019; Ocasio et al., 2023.

### SCENIC

The SCENIC algorithm (Single-Cell Regulatory Network Inference and Clustering) reconstructs regulons—transcription factors (TFs) and their target genes—from scRNA-seq data. It involves three main steps: (1) regulatory network inference using GRNBoost2, which identifies TF-gene coexpression patterns through a gradient boosting machine algorithm; (2) pruning indirect targets via cisTarget, leveraging motif enrichment and cis-regulatory module analysis to refine regulons; and (3) quantifying regulon activity across cells using AUCell, which scores regulon target gene enrichment with an area under the curve (AUC) metric. Key hyperparameters include the number of tree estimators in GRNBoost2, motif enrichment thresholds in cisTarget, and regulon AUC thresholds. The SCENIC workflow identified TFs enriched in regulons, demonstrating their role in cellular regulatory networks (Van de Sande et al., 2020).

### RBM

The Restricted Boltzmann Machine (RBM) is implemented using the scikit-learn library (BernoulliRBM) with key parameters including n_components=100 (hidden units), learning_rate=0.01, batch_size=10, n_iter=25, and random_state=42. RBM inferred attractors by fitting log-normalized single-cell expression data and generating latent samples via Markov Chain Monte Carlo (MCMC) sampling. Dimensionality reduction (t-SNE, UMAP) visualized latent spaces of the RBM trajectories, while K-means clustering (optimized between 2-10 clusters) identified optimal attractors using silhouette scores. Differentially expressed genes (DEGs) were detected within clusters using two-sample t-tests (p < 0.05), revealing genes critical for each cluster’s identity. RBM mapping predicts transcriptional features underlying cellular heterogeneity. Random Forest identified the top 100 genes. (See Data and Code Availability).

### Network Analysis

The Partial Information Decomposition (PID) framework is a computational method used to analyze gene regulatory networks by decomposing the information shared between variables into unique, redundant, and synergistic components. Utilizing Julia’s NetworkInference.jl package, the method takes gene expression data (e.g., K27M and IDHWT datasets) as input in text file format. The PIDC algorithm infers dependencies between genes by breaking down mutual information into pairwise contributions, constructing networks where nodes represent genes and edges capture regulatory relationships. The graph R package computed centrality measures including eigenvector centrality, betweenness centrality, and closeness centrality, on the undirected graphs. Spearman adjacency matrix was also used for the histone-histone post-translational modification (PTM) networks. Mann-Whitney U test and one-sample t-tests were used to assess the statistical significance of observed values (e.g., centrality measures, BDM changes, importance scores).

### Histone Networks

Histone-histone co-modification networks were generated from the histone modification profiling data from the Cancer Cell Line Encyclopedia (CCLE) glioma samples (Ghandi et al., 2019) and the BT245 cell line (Harpaz et al., 2022). Histone mass spectrometry and Cytometry by Time of Flight (CyTOF) were used to examine a broad panel of PTMs in these datasets, respectively. For the DIPG BT245 samples (Harpaz et al., 2022), CyTOF data for a wide array of histone modifications at the single-cell level, were obtained from a custom-designed epigenetic panel. The relative abundance matrix represents the proportion of specific histone modifications, calculated as the ratio of the area of a modification (e.g., H3K4me3) to the total area for all modifications of the same peptide (e.g., H3_3_8). These ratios indicate the prevalence of each histone modification. The CCLE data was median-normalized, log-transformed, and comparable to z-scores for consistency and interpretability. Spearman correlation and PID were used to construct adjacency matrices. The resulting networks underwent BDM perturbation analysis, and centrality measures analysis.

### Block Decomposition Method (BDM)

The BDM is a measure of algorithmic complexity, quantifying the informational content of a system based on the estimated Kolmogorov complexity (akin to compressibility) of its adjacency matrix (Soler-Toscano et al., 2014; Zenil et al., 2017). BDM perturbation analysis involved binarizing the network adjacency matrix with a threshold of 0.5 for the log-normalized counts and assessing BDM shifts in network topology using the pyBDM python package. The igraph R package and network python packages were used for graph construction and centrality computations, and shifts in key nodes were ranked to identify the top-affected genes.

### Langevin Dynamics

Cell fate trajectories were stimulated by their gene expression dynamics using Ito stochastic differential equations (SDEs) within the Langevin framework, where the drift term models deterministic forces, and the noise term represents stochastic perturbations. Initial parameters (α=1.0,β=0.5,γ=0.1=1.0) were optimized using L-BFGS-B and differential evolution (DE) algorithms to minimize the mean squared error (MSE) between simulated and actual gene expression data. Simulations were performed over T=10 units of time with dt=0.01 time steps, resulting in 1000 simulations per gene. Clustering (K-means, k=3k) and PCA reduced the dimensionality of simulated trajectories, while a Random Forest classifier identified key genes by feature importance, enabling insights into cell fate dynamics. Packages include Scipy, Scikit-learn, and Matplotlib for optimization, clustering, and visualization. The optimized parameters from the evolutionary algorithm for K27M were [0.1005,0.1008,0.0109][0.1005, 0.1008, 0.0109][0.1005,0.1008,0.0109], and for IDHWT were [7.2599,1.9612,0.0146][7.2599, 1.9612, 0.0146][7.2599,1.9612,0.0146].

### Bayesian Inference

The BASiCS algorithm (R package) performs Bayesian network-based causal inference on single-cell RNA-seq data to estimate gene-specific parameters using Markov Chain Monte Carlo (MCMC) (Vallejos et al., 2015). Cells were split into two random batches, and posterior distributions were inferred for key parameters: μ (mean gene expression), δ (over-dispersion indicating variability beyond Poisson noise), and σ (total variability). Highly variable genes (HVGs) are identified by thresholding δ, with higher δ values indicating significant expression variability. The algorithm effectively distinguishes HVGs and lowly variable genes (LVGs) to reveal important transcriptional heterogeneity across cells. Parameters were optimized with N=100N=100N=100, thinning = 5, and burn-in = 50 iterations.

### Gene Ontology and Gene Set Enrichment Analysis

The g:Profiler tool, gene set enrichment analysis was performed to identify biological processes, pathways, and functional categories associated with a list of gene markers (Raudvere et al., 2021). Additionally, the GeneCards database (Stelzer et al., 2016) was utilized to validate and explore further the functional roles and interactions of these genes, mentioned in the Discussion Section.

## RESULTS

### scEpath Waddington landscape reconstruction identifies histone demethylases, and developmental genes as pHGG cell fate dynamics transition markers

Various transcription factors were identified as transition markers in pediatric K27M DIPG and IDHWT glioblastoma, with statistically significant cell fate trajectories mapped onto a Waddington energy landscape. K27M glioma exhibited three differentiation paths, while IDHWT glioblastoma showed two bifurcating cell fate trajectories, with high-energy states corresponding to glioma stem cells. Transition genes shared across both systems include KDM5B, TGIF1, SOX11, and HMGB3. Lineage-specific markers such as OLIG1, SOX10, and SOX6 in K27M glioma and SOX11 in glioblastoma underscore divergent differentiation dynamics. SOX4, a paralog of SOX11 (P = 1.08E-06), emerged as a critical cell fate transition gene in K27M, while BDM perturbation analysis revealed significant interactions such as ETV5-ID3 (P = 6.95E-06) and TGIF1-TSC22D4 (P = 6.95E-06). In pediatric glioblastoma, the ARID5B-KDM5B interaction demonstrated the highest BDM perturbation shift (8.42 bits), highlighting their central roles in epigenetic regulation through synergistic H3K4 and H3K9 methylation patterns. Other key interactions included ARID5B-HMGB3 and DLX1-KDM5B, with significant shifts of 4.95 bits each. Genes such as DLX5 and DLX6, involved in brain patterning and neuron-glial integration, were highly ranked in centrality measures. Central regulators with the highest eigenvector centrality in K27M included SATB1, ARNT2, and TGIF1, while KDM5B (H3K4me2/3 demethylase) and ARID5B (H3K9Me2 demethylase) were identified as key network hubs in IDHWT glioblastoma (P = 4.62E-25).

**Figure 1.**
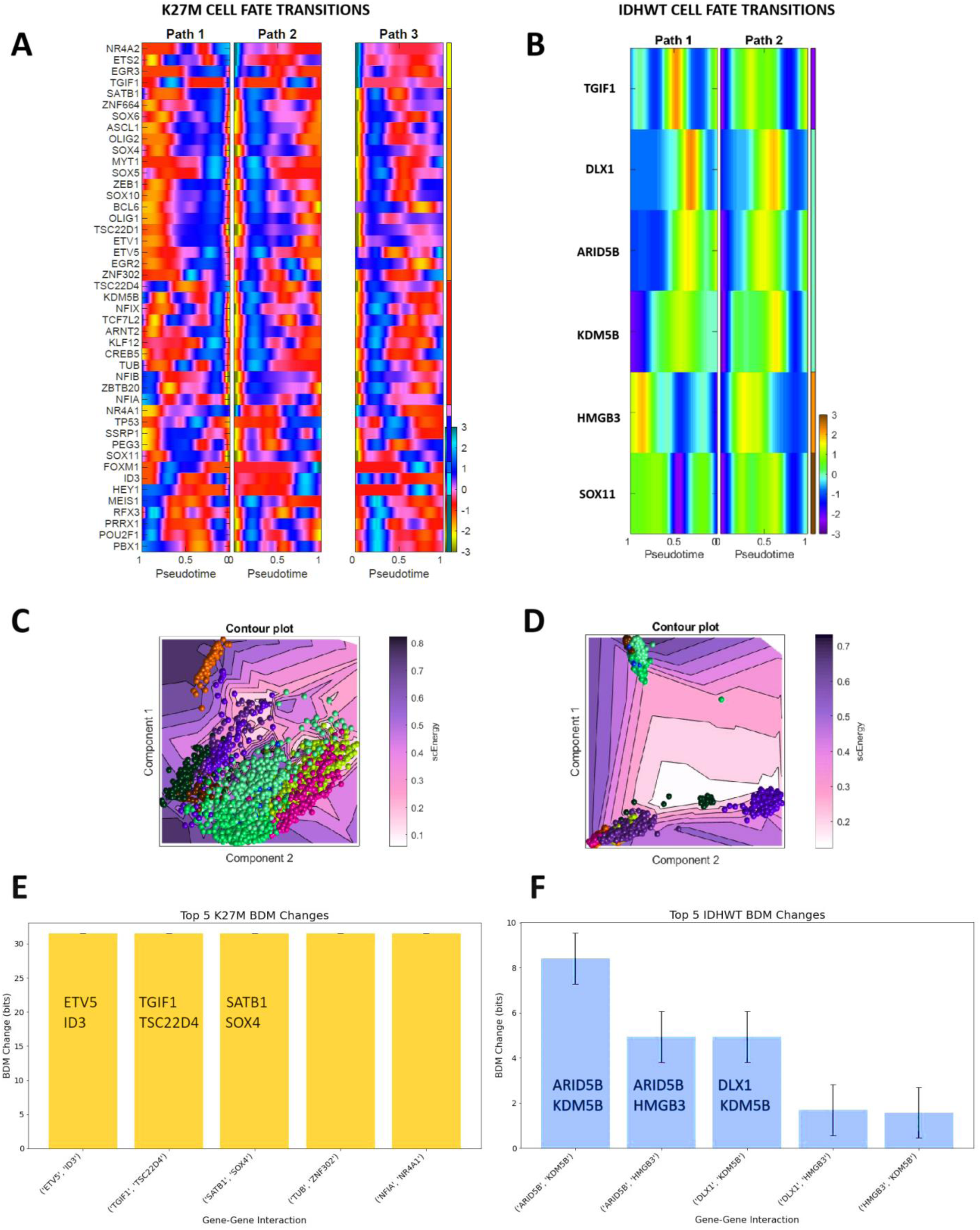
scEpath Waddington energy landscape reconstruction infers attractor dynamics and histone demethylases as transition markers in glioblastoma cell fate dynamics. A) Three bifurcation paths from high energy cell states with transition genes for K27M glioma shown in pseudotemporal progression. B) Two bifurcation paths for cell fate transitions in IDHWT glioblastoma with key differentiation genes. The normalized gene expression gradient is shown in the color bar. C) Contour plot of the Waddington-like energy landscape for K27M glioma. The color bar denotes the normalized gene expression level during the fate transitions/trajectories. D) Contour plot of energy landscape for IDHWT glioblastoma. E) Algorithmic complexity perturbation analysis of top transition gene-gene interactions in K27M. F) Algorithmic complexity perturbation analysis of highest transition gene-gene interactions in IDHWT. Many of these markers are critical for brain patterning and neurodevelopmental regulation.

### SCENIC and CellRouter Algorithms Highlight Regulatory Genes Orchestrating Developmental Processes as Candidate Biomarkers of pHGG Phenotypic Transitions in Lineage-specific patterning

As shown in Figures 2A and 2B, for K27M glioma, the key regulons identified included ETV2, IKZF1, IRF8, KLF14, and MYC, with enrichment pointing to roles in myeloid differentiation and hematopoiesis. Other significant regulons, such as ELK3, FOXO3, and FOZG1, were associated with glioblastoma plasticity transitions, including SMAD3/4-FOXO3-FOXG1 complex formation and mitotic activity. Top BDM perturbation shifts highlighted interactions such as ETV3-FOXO3 and ATF3-E2F7, emphasizing FOXO3’s involvement in oxidative stress, T-cell regulation, and apoptosis, and ETV3’s role in KRAS-regulated gene systems. These findings suggest that developmental signatures and immune-inflammatory processes linked to oxidative stress are critical in K27M glioma transcriptional dynamics. For IDHWT glioblastoma, the top regulons included GATA2, ETV7, HOXB6, ARX, and FOXN1, with enrichment indicating roles in organ morphogenesis and neuronal differentiation. Key BDM shifts, such as FOXA2-HES7 and ATF3-E2F1, highlight FOXA2’s role in early developmental processes and HES7’s involvement in somitogenesis and tissue patterning (Bessho et al., 2001; Liu et al., 2024). ATF3-E2F interaction, observed in both gliomas, serves as a potential biomarker, with E2F7 specific to K27M and E2F1 to IDHWT. Both E2F genes interact with EZH2, linking PRC2-mediated chromatin regulation to glioblastoma progression through distinct pathways (Yang et al., 2020). As shown in Figure 3, ATF3 interacts with JUN, which has been validated in leukemias (Zhou et al., 2017).

**Figure 2.**
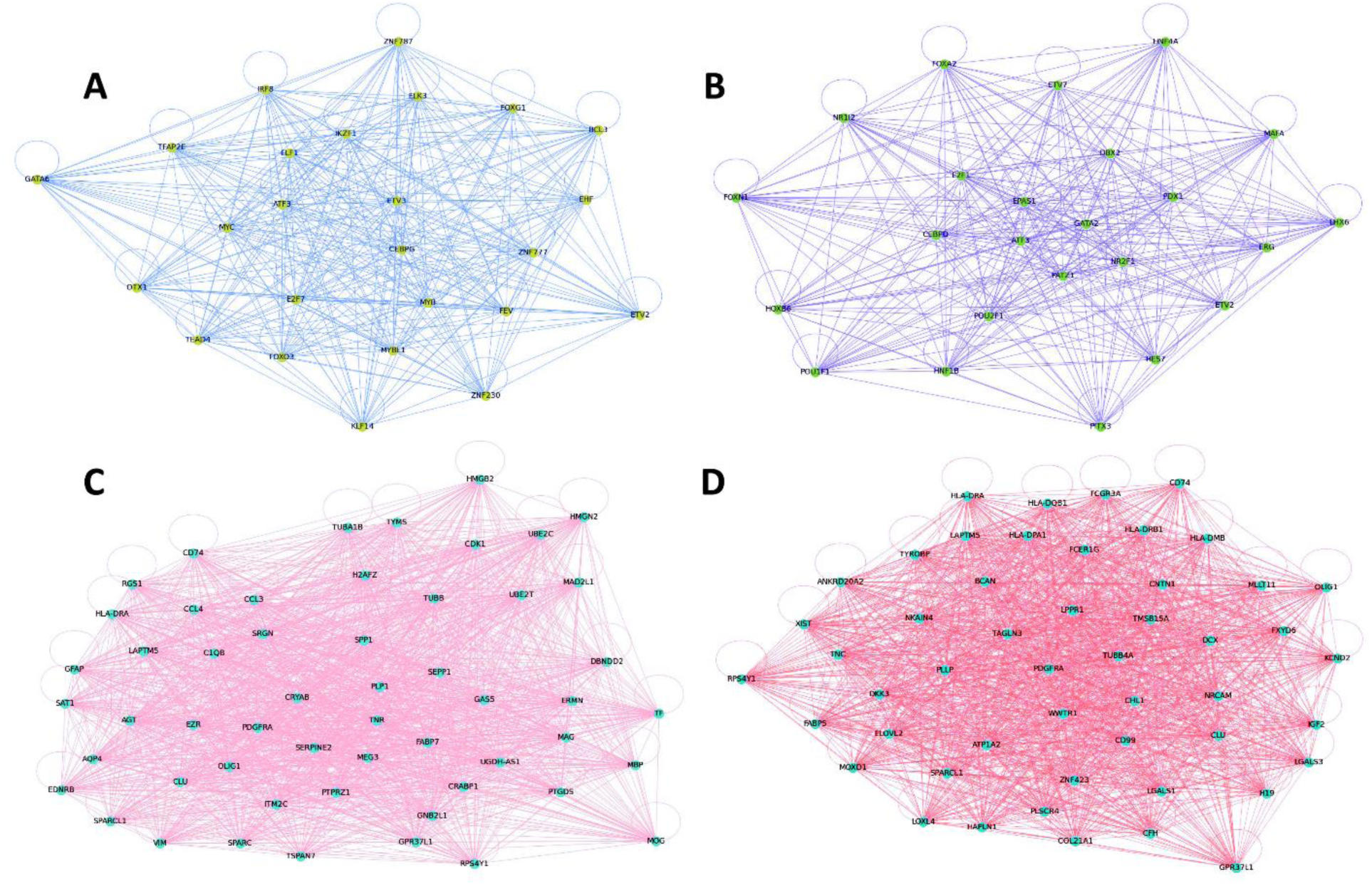
Network science identifies central regulons and molecular features of cell fate decisions in single-cell trajectory inference algorithms. A) Regulon networks using Spearman correlation from SCENIC algorithm for K27M. B) Regulon networks from SCENIC for IDHWT. C) Top molecular features in Louvain community clusters across K27M inferred by CellRouter algorithm. D) Top Louvain community cluster markers in IDHWT by CellRouter.

**Figure 3.**
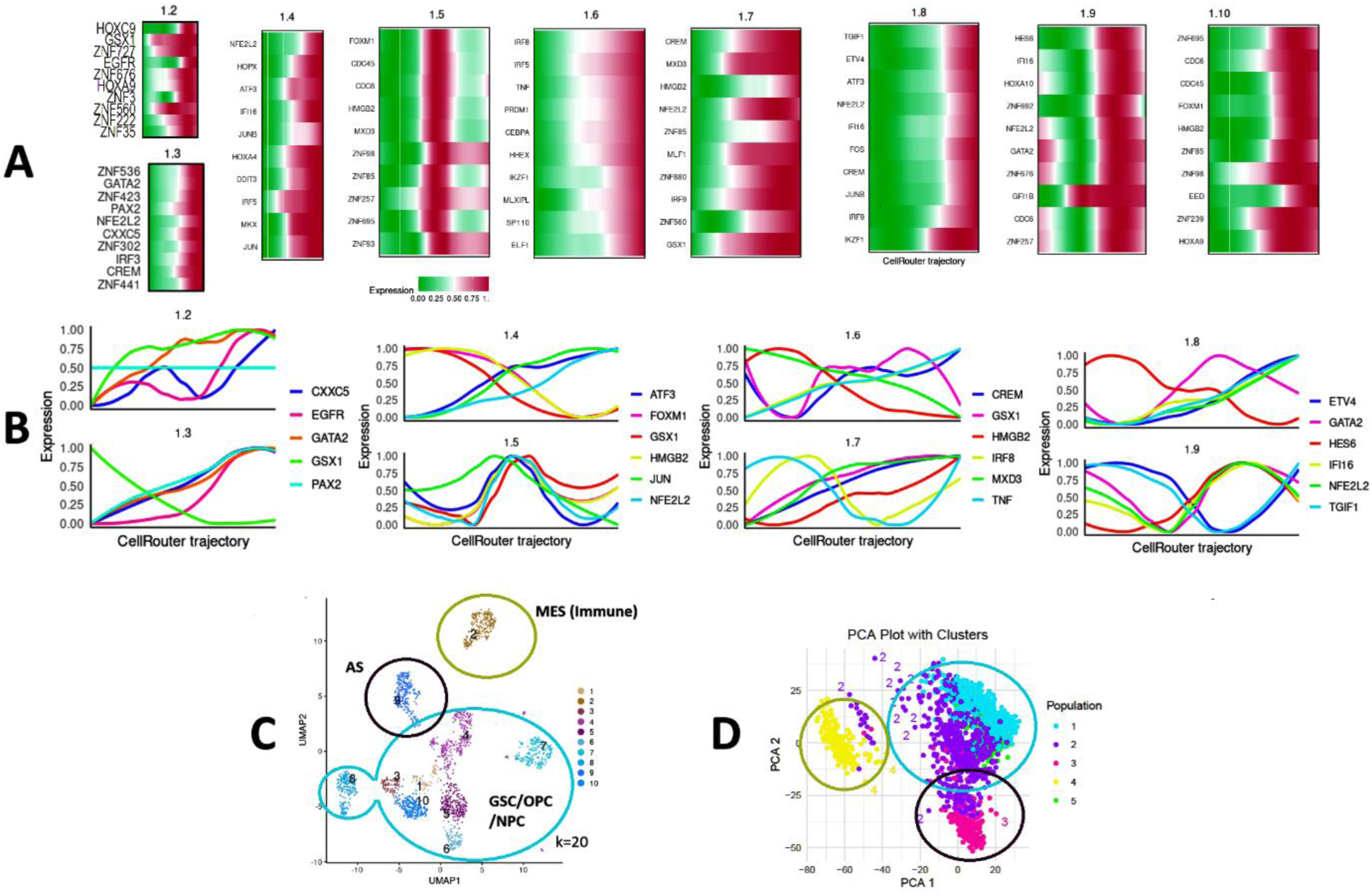
CellRouter Trajectory Inference Algorithm Identifies Kinetic Dynamics and Complex Oscillations of Key Transcription Factors Steering Cell Fate Decisions in IDHWT Glioblastoma (k=20). A) State-transition genes inferred by CellRouter for differentiation trajectories, with transitions from Louvain community cluster 1 to the cluster denoted after the decimal. The color scale shows normalized gene expression from green (0.0; low) to red (1.0; high) in pseudotime. B) Kinetic curves of transcription factors driving differentiation trajectories. C) UMAP embedding of glioblastoma cell states clustered by kNN graph network reveals a trilineage of potential differentiation trajectories. D) PCA dimensionality reduction of kNN graph clustering, showing transcriptional cell-state populations: The neural stem cells (NSCs) and their bifurcations into GSC/OPC/NPC hybrid states (light blue), astrocyte-like states (black), and mesenchymal-like states (green). Tumor ecosystems exhibit collective intelligence driven by spatiotemporal gene expression and protein distribution, guiding phenotypic plasticity transitions.

In K27M, FOXO1 emerged as a key feature, while in IDHWT, FOXD4L4 and MAGED4 showed the highest importance scores. MAGED4, shared across both gliomas, plays a role in ubiquitin ligase activity, potentially influencing histone network dynamics and chromatin remodeling. These findings, with BDM shifts exceeding 30 bits, highlight critical transcriptional regulators and epigenetic control mechanisms driving glioma plasticity and offer insights into lineage-specific therapeutic targets.

### CellRouter Trajectory Inference Algorithm Predicts the Cellular Identities of IDHWT Glioblastoma and Identifies Hematopoietic/Immune Progenitor States Coexisting with Neural Stem Cells

Using CellRouter as an unsupervised trajectory inference algorithm, Figure 3A shows the various transition pathways in UMAP or tSNE embedded feature space with k=20 clusters in the graph network by similarity in gene expression. We identified ten Louvain community clusters representing diverse phenotypic subpopulations within glioblastoma. Key transition markers and pathways revealed complex lineage plasticity dynamics. Panel 1.2 indicates mesenchymal stem/progenitor cells characterized by HOXC9, HOXA9, and EGFR, highlighting their roles in epithelial-mesenchymal transitions (EMT) and mesenchymal differentiation. Panel 1.3 features neural progenitors with markers like GATA2, PAX2, and NFE2L2, suggesting a neural progenitor state connected to hematopoietic lineages and early embryonic somitogenesis. Panel 1.4 identifies stress-responsive progenitor cells expressing NFE2L2, HOPX, and ATF3, while Panel 1.5 represents glioma stem cells (GSCs) marked by FOXM1, CDC45, and HMGB2, denoting highly proliferative states. Panel 1.6 characterizes mesenchymal (stem) cells with overlapping hematopoietic and immune progenitor cell markers, such as IRF8, CEBPA, and TNF, indicating immune-inflammatory responses from mesenchymal or hematopoietic lineage commitment. Panels 1.7–1.9 represent NPCs (neural progenitor cells), astrocyte precursors, and differentiated astrocytes, respectively, with markers like TGIF1, HES6, and HOXA10, while Panel 1.10 captures proliferative GSCs expressing ZNF695, FOXM1, and CDC6. For instance, HES6 is critical for the switch between astrocytic and neuronal differentiation (Jhas et al., 2006).

Distinct transcriptional modules validated previous findings on glioma lineage origins. Markers such as PDGFRA, OLIG1, and SOX10 emerged in OPC-related modules, while GATA2 and HOXA9 linked neural and hematopoietic lineages, emphasizing shared plasticity across these populations (Mendez-Gonzalez et al., 2019; Larsson et al., 2024). NFE2L2, a regulator of oxidative stress, was prevalent across all clusters, underscoring its role in sustaining metabolic adaptability. Oscillatory gene expression patterns, such as ATF3, GSX1, and IRF8, suggest collective oscillatory dynamics driving glioblastoma transitions towards immune-mesenchymal states. Tumor ecosystems exhibit emergent nonlinear systems and decentralized control, forming a swarm intelligence that integrates multiscale gene-environment interactions. This continuum of adaptive stem cell states, encompassing NSC, GSC, NPC, OPC, and HSC markers, reinforces glioblastoma as a *complex adaptive system* with high phenotypic plasticity.

### Restricted Boltzmann Machines (RBM) and Bayesian Network Inference reveal Oncofetal Signatures, Telomere Lengthening, and Developmental Biomarkers as Drivers of Plasticity in pHGG cybernetics

Figures 4A and 4B demonstrate the RBM-inferred differentiation trajectories from neural stem cells (NSCs) and glioma stem cells (GSCs) to differentiated states such as mesenchymal progenitors. RBMs predicted causal evolutionary dynamics across lineage bifurcations, with key genes in K27M including SPATA2 (P=4.75E-05), DRAXIN (P=5.27E-13), and MTSS1 (P=6.82E-10). These genes are associated with TNF signaling, axon guidance, EMT transitions, and developmental plasticity. Additional features such as CAMK2D (P=5.12E-13) and FGFR1 (P=4.37E-06) highlight roles in calcium dynamics, synaptic plasticity, and cell signaling. Wnt, Frizzled, and SMAD/STAT3 signaling pathways were enriched, reflecting hijacked oncofetal developmental circuits. Oncofetal developmental networks refer to the reactivation of embryonic (neurodevelopmental) gene expression programs in tumors, driving phenotypic plasticity, stemness, and malignancy due to disrupted development processes. Meanwhile, IDHWT glioblastoma showed top features like DKK3, SLC3A2, and RLIM (P=0.00014), the latter being involved in telomere length regulation and immortalization. Bioelectric markers such as GRIK3 and GRIN3B, ligand-gated ion channels, underscore the role of endogenous voltage-gated networks in phenotypic plasticity and EMT transitions.

### Bioelectric and Immune-Proinflammatory Signatures

Glioblastoma revealed critical roles for bioelectric and proinflammatory markers within its top 100 most important features predicted by RBM. GRIK3, linked to EMT transitions and cancer invasion, and TOB1, involved in BMP signaling and EGFR pathways, were observed in the IDHWT glioblastoma. Shared features across both glioma systems, such as RLIM and TERF1, emphasize the hijacking of telomere lengthening processes. Tumor-associated macrophage (TAM) signatures like C1QA and CD59 highlight the neuroendocrine regulation of immune signaling. Cytoskeletal regulators, including CDC42EP5 and CTNND1, highlight ECM-cytoskeletal dynamics in glioma plasticity. Collectively, these findings demonstrate that pHGG ecosystems reprogram bioelectric and proinflammatory networks to drive plasticity and phenotypic transitions through complex, multiscale cybernetics (networks).

**Figure 4.**
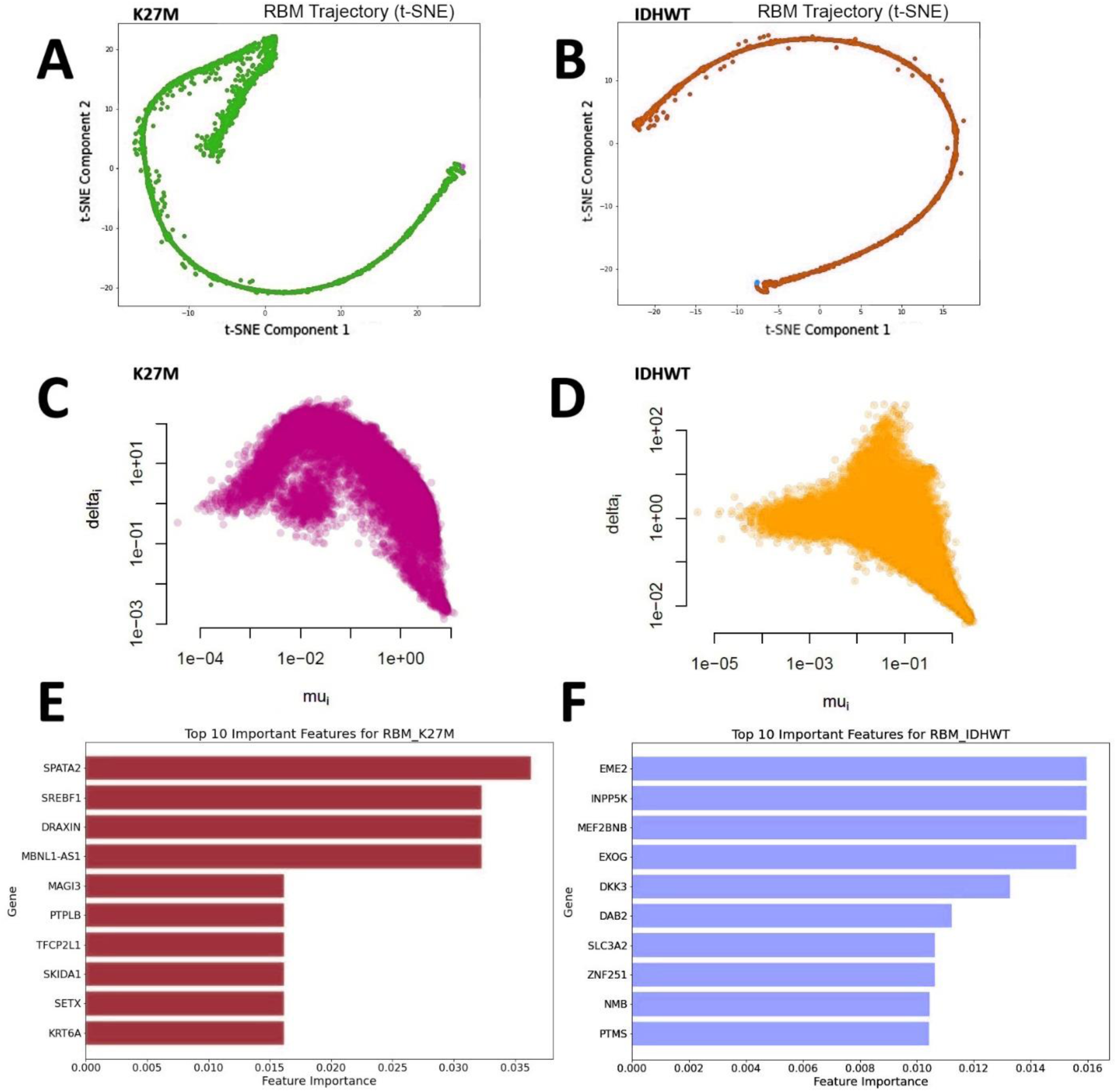
Bernoulli Restricted Boltzmann Machines (RBM) and Bayesian Network Inference Predict Transition Signatures in pHGG gene expression. A) RBM trajectory inference of K27M glioma cells by gene expression in t-SNE feature space. B) RBM cell fate trajectory inference on IDHWT glioblastoma in t-SNE space. C) Bayesian network inference of K27M cell fate choices. D) Bayesian network inference of IDHWT cell fate dynamics. The parameter Mu represents the mean expression level of the gene across the cells. A trilineage attractor of three differentiation pathways (NPC/neuron-like, OPC/astrocyte-like, and mesenchymal/microglia-like) can be seen in both pHGG systems. The parameter Delta **δ** represents the over-dispersion of the gene’s expression levels. E) Top 10 important features of K27M glioma extracted using random forests (RF) on RBM signatures. F) Top 10 important features of IDHWT glioblastoma in RBM decision space. Importance features, or salience maps, provide explainable AI methods for biomarker discovery.

### Partial Information Decomposition (PID) Networks of RBM and Bayesian Inference-based Importance Features Reveal Cytoskeletal Remodelling, Exosome-mediated Vesicular Transport, and Immune-Inflammatory Signals as Regulators of Glioma Progression

Bayesian network analysis of K27M glioma revealed significant immune-inflammatory responses, with key HVGs such as CCL3 (sigma = 0.964, P = 2.15E-131), CCL4, and TYROBP. Microglial markers C1QC, C1QB, and FCER1G were enriched, suggesting active immune cell dynamics and microglial activation. The top gene-gene interactions in K27M, identified by BDM perturbation shifts, included CX3CR1-SELL, APBB1IP-MIR766, and CH25H-IL1B (P < 1E-132). These interactions highlight microglia-specific immune signals as potential immunotherapy targets. Among HVGs, CD74, HLA-DRB1, and CYBB showed the highest eigenvector centralities (>0.9), indicating strong hub roles. LAPTM5, a centrality gene in K27M dynamics, emerged as a microglial signature and precision therapy target for inhibiting invasion and chemoresistance (Berberich et al., 2020). In IDHWT glioblastoma, immune responses were characterized by C1QA, HLA-DRA1, and SNAR-A10. Key HVG interactions included C3-SNORD115-4 (P = 1.20E-17) and CCL3-SNAR-A11 (P = 1.14E-16), implicating the complement system and inflammatory signals in glioma plasticity dynamics. Central genes TYROBP, LAPTM5, and VSIG4 showed high eigenvector centrality (1.0) and betweenness centrality (690.33), confirming their roles in hematopoietic differentiation and immune regulation. Immune progenitors within tumor ecosystems interact with CSCs, creating a decentralized tumor-immune landscape (TME) that coordinates tumor morphogenesis and neuroendocrine-immune signaling.

In the PID network for K27M glioma, the highest BDM shifts included VTA1-XPO6, GABARAPL1-HFM1, and BIRC6-C6orf123. VTA1 (C6orf55) and XPO6 are involved in vesicular trafficking and actin dynamics, with links to EGFR signaling. GABARAPL1 plays a critical role in mitochondrial processes, autophagy, and senescence, highlighting the involvement of oxidative stress in tumor dynamics. BIRC6, a ubiquitin ligase, regulates apoptosis and cellular homeostasis. These interactions emphasize mitochondrial activity, lysosomal degradation, and ATP-dependent cellular processes as pivotal in glioblastoma plasticity. In IDHWT glioma, key BDM shifts included SLC5A9-SMARCAL1, CD59-CLEC11A, and KIAA0753-TUBB1. SLC5A9, a sodium ion transporter found on exosomes, suggests potential for liquid biopsy diagnostics. SMARCAL1, a chromatin remodeling gene, and CLEC11A, a growth factor for HSC maintenance, further validate the link between HSC and neural stem cell (NSC) plasticity. TUBB1, involved in microtubule polymerization, and KIAA0753, critical for centrosome regulation, underscore the role of microtubule-mediated processes in tumor evolution. These findings demonstrate how mitochondrial oxidative stress, stem cell dynamics, and chromatin remodeling collectively drive glioblastoma plasticity and offer targets for therapeutic intervention.

As shown in Table 5, C6orf123 appeared among both the highest positive and negative BDM shifts in K27M, highlighting its dual role in glioblastoma dynamics. DOCK7 (P=6.92E-10), involved in axon formation and radial glia migration, underscores Rho GTPase signaling as a potential target for reprogramming glioma plasticity. The KCNJ6-ATG3 interaction (P=1.20E-06 for both genes) emerged as a significant endogenous bioelectric network, with KCNJ6 encoding a potassium ion channel that could modulate phenotypic plasticity. HHEX, a hematopoietic transcription factor, and WNT activator, was linked to CTNNB1 (β-catenin), identified as a key proteomic marker in pediatric gliomas (Petralia et al., 2020). HHEX was also identified in IDHWT glioblastoma as a transition gene by the CellRouter algorithm (Figure 3). This reinforces the role of WNT/β-catenin signaling in glioblastoma progression (Wang et al., 2021). TGFBI (P=9.65E-22) further highlighted the importance of collagen and extracellular matrix remodeling via TGF-beta pathways.

In IDHWT, DKK3, a WNT-associated gene, showed the highest negative BDM shift. NOTCH2NL, involved in cortical plasticity, and TRPM6, an ion channel gene, emerged as significant RBM markers. ANXA1-IL1B interaction (P=1.36E-19) was the most negative BDM shift in K27M, where ANXA1 promotes tumor immune evasion and IL1B regulates macrophage responses, making it a major immunotherapy target. Similarly, APOC1-ADORA3 interaction highlighted synergetic immune cell regulation roles. In IDHWT, GAGE8 (P=4.13E-16), aberrantly expressed in tumors, indicated potential immune functions. These findings collectively identify WNT signaling as a shared vulnerability across glioma systems and a target for phenotypic reprogramming. Bioelectric networks (e.g., KCNJ6), Wnt, and Notch-based signaling networks, and microenvironmental immune-inflammatory interactions (e.g., ANXA1-IL1B) offer additional precision therapeutic avenues for controlling glioma cell plasticity and progression.

**Figure 5.**
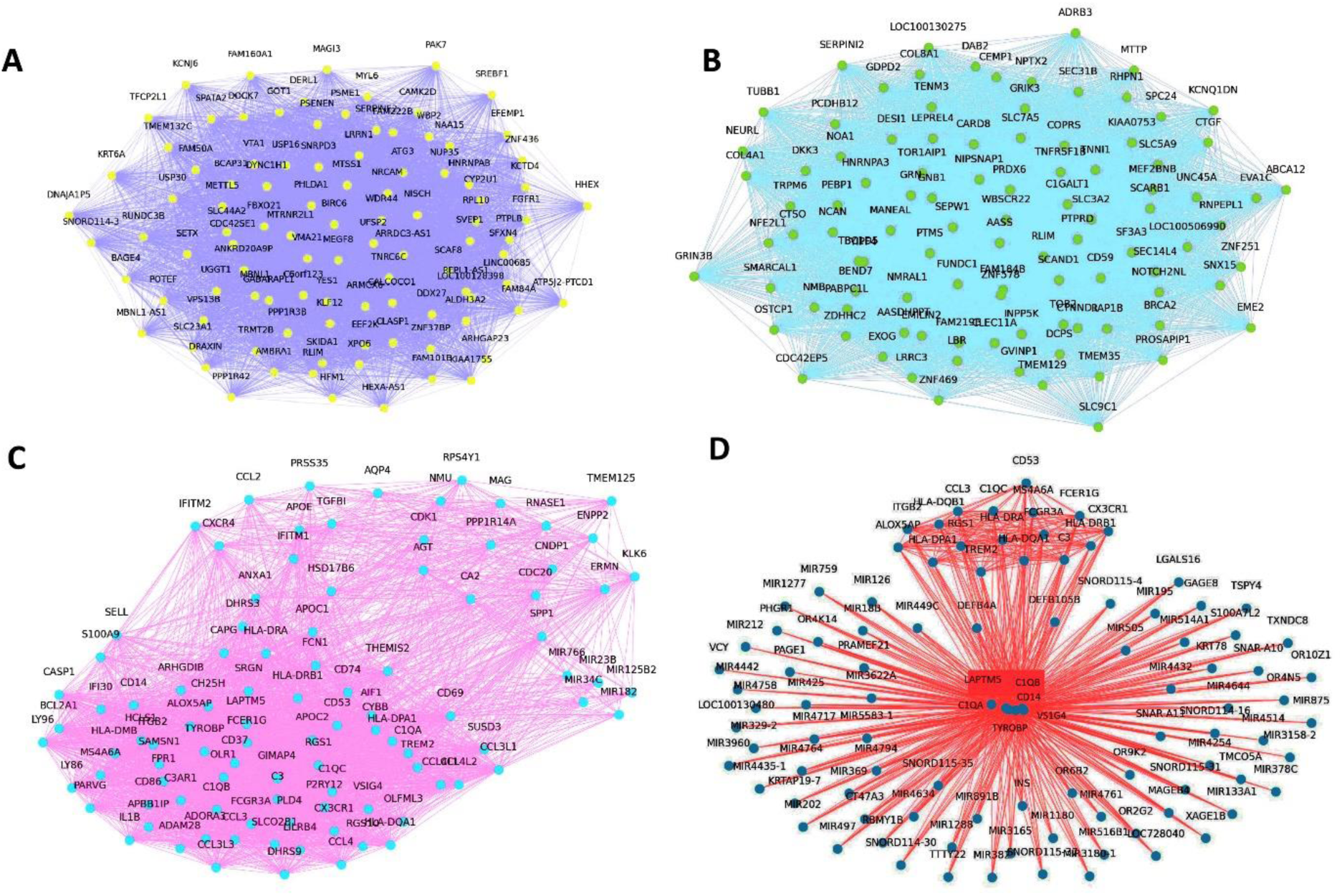
Network Dynamics of HVGs and Importance Features Extracted by Bayesian Network Inference and RBM probabilistic graphical models using the PID metric. A) BASIC algorithm inferred Bayesian Network of Important Features in K27M. B) Bayesian Network Inference in IDHWT glioblastoma. C) RBM inferred transcriptional networks of important features in K27M. D) RBM inferred networks in IDHWT.

### Bayesian Network Dynamics Reveal Signatures of Oxidative Stress Metabolism

In the Bayesian HVGs inferred by the BASIC algorithm, 21 shared genes between K27M and IDHWT included microglial signatures involved in oxidative stress and vesicle-mediated communication. C1QA, C1QB (Mann-Whitney p=6.38E-14), HLA-DRB1 (p=1.47E-05), and ITGB2 were linked to elevated C-reactive protein (CRP), suggesting potential liquid biopsy biomarkers. GATAD2A, part of the NuRD complex, emerged as a central regulator of these shared genes, emphasizing its epigenetic role in glioblastoma dynamics.

For K27M, MIR23B and MIR182 showed the highest centralities (betweenness = 271.98, eigenvector = 1; p=3.01E-07). MIR23B downregulates IGF1 via GATA6, observed in SCENIC regulons, while MIR182 has known glioblastoma associations. In IDHWT, C1QA and C1QB had the highest centralities (betweenness ∼40; eigenvector = 1), with ALOX5AP (p=1.46E-17), encoding arachidonic acid metabolites, ranked third. FABP5, a regulator of endocannabinoid transport linked to arachidonic acid, was previously identified in K27M. Centrality measures for K27M PID networks were statistically significant (p-values <1E-12).

### RBM Network Dynamics Identify ROS Metabolism and Differentiation Markers

In K27M, RBM analysis revealed LRRN1 as the highest betweenness (85.84) and MBNL1 as the highest eigenvector centrality (0.96; p=5.30E-65), with roles in synaptic function, ROS metabolism, and alternative splicing. In IDHWT, RAP1B exhibited the highest centralities (betweenness = 58.97, eigenvector = 1), highlighting its role in focal adhesion dynamics and differentiation. SF3A3 and MANEAL, involved in splicing and ECM remodeling, also emerged as top features. These findings underscore the shared importance of ROS regulation, endocannabinoid signaling, and epigenetic control across glioma systems, providing actionable targets for precision therapy.

### Stochastic Modelling with Langevin Dynamics Identifies Similar Epigenetic Signatures Steering pHGG Cell Fate Decisions and Glioma Plasticity

Langevin Dynamics, optimized by Evolutionary Algorithms, modeled transcriptional state transitions in glioma cell fate trajectories (k=3 clusters), complementing previous algorithms. As shown in Figures 6A and 6B, the Langevin simulation coupled with the EA is shown to infer the cell fate trajectories by each gene across time steps until it settles to equilibrium states. Figures 6C and 6D show the transition dynamics between three k-means clusters (k=3). The cell fate transition trajectories are smoother in K27M (Fig 6C) than in IDHWT (Fig 6D) suggesting a better-optimized model fit. Figures 6E and 6F reveal the random Forest (RF) extracted importance features (gene signatures) from the inferred cell fate trajectories. BDM perturbation analysis on single-gene signatures was performed to identify information signatures regulating the cell fate choices.

**Figure 6.**
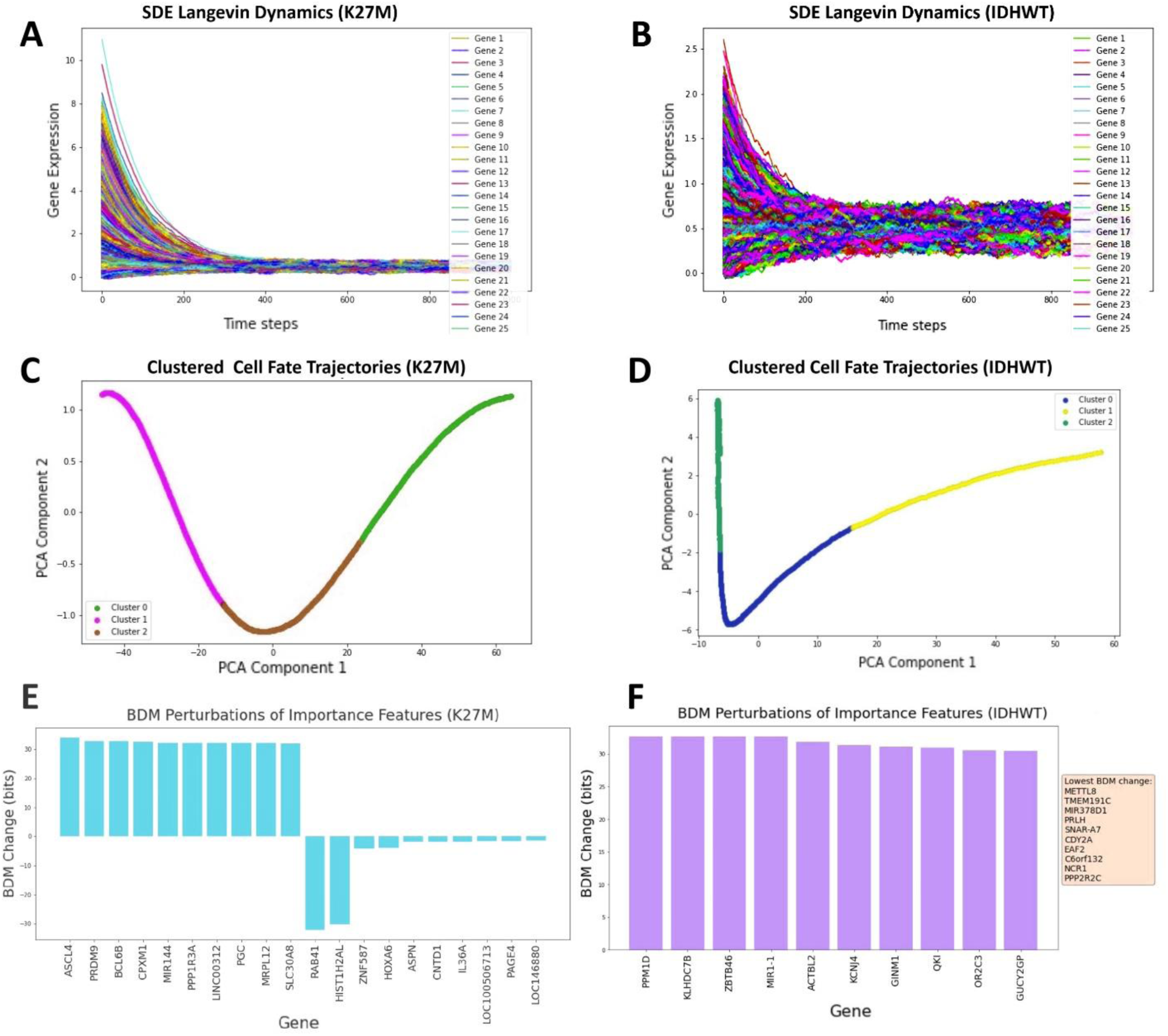
Langevin Dynamics simulations using Stochastic Differential Equations (SDE) Modelling show Markovian processes can capture inferred attractor network dynamics in pHGG systems. A-F) Langevin simulations on K27M (A) and IDHWT (B) gliomas modeled cell fate trajectories, visualized in PCA space clustered using k-means for K27M (C) and IDHWT (D). E) Algorithmic complexity (BDM) perturbation in K27M identified importance genes with Mann-Whitney p=7.92E-07. F) In IDHWT, perturbation signatures had p=0.026. These findings suggest potential applications for modeling multiscale pattern formation via bifurcations, reaction-diffusion systems, and active turbulence in tumor dynamics.

Figures 6A-F highlight Langevin dynamics-inferred cell fate trajectories and gene transitions across K27M and IDHWT gliomas. In K27M, the highest BDM shifts (>30 bits) were observed for ASCL4, PRDM9, BLCGB, and CPMX1. ASCL4 is linked to glioma metabolism and proliferation, while PRDM9 catalyzes H3K4me3, and is critical for epigenetic regulation (Diagouraga et al., 2018; GeneCard Database).

Telomeric markers, including TERF1, were identified, further validating the importance of telomere regulation, supported by RLIM-mediated inhibition of TERF1. Other key signatures include SEZ6L1 and HOXA6, associated with NSC and nervous system development, and migrasome-related genes TOMM20 and HSPD1 (as identified by g:Profiler analysis), which underscore the role of exosomal communication in glioma progression (Zhang et al., 2023).

In IDHWT, PPM1D (a p53 regulator) had the highest BDM perturbation, followed by KLHDC7B and ZBT46. Epigenetic and bioelectric markers, including METTL8 (mRNA methyltransferase), CDY2A (histone acetyltransferase), and GRIA3 (AMPA receptor gene), emerged as critical regulators. The KCNJ4/6 potassium channels were highlighted as strong candidates for bioelectric reprogramming of glioma phenotypes. Both glioma systems shared TGM6, a transglutaminase linked to leukemia. Methylation dynamics, including H3K4me3 (PRDM9), bioelectric ion channels (KCNJ4/6), and migrasome-mediated signaling, were central to glioma progression and resilience. These findings identify actionable targets for epigenetic and bioelectric reprogramming to mitigate phenotypic plasticity and tumor aggressiveness, providing potential avenues for precision therapies. Again, most of these functional roles are extracted from the GeneCard Database and g:Profiler analysis.

### Single-cell Mass Cytometry Reveals H3K4 and H3K9 Methylation Signatures as Central Drivers of Histone-Histone PTM Networks

Figures 7A and 7B show PID networks of histone modifications in gliomas from the CCLE database and the BT245 K27M glioma cell line, respectively, while Figure 7C illustrates the CCLE Bayesian network clustered by Louvain detection. Centrality measures reveal H3K4me1 as the highest betweenness and closeness centrality in both CCLE PID and Bayesian networks. The highest eigenvector centrality in the CCLE networks was H3K27ac1K36me2, while H3K9me2K14ac0 showed the highest closeness in the Bayesian network. These findings suggest that H3K4 and H3K9 modifications dominate transitions between stem cell-like and differentiated states, while H3K36 methylation serves as a regulatory switch. In the BT245 histone network, H3K4me3 had the highest eigenvector centrality, while H3K9me3 showed the highest betweenness and closeness centralities, followed by H3K27me3, a hallmark of K27M gliomas. The second-highest eigenvector centrality was H4K16ac, further emphasizing the importance of H3K4 and H3K9 methylation in epigenetic plasticity. These consistent patterns across PID and Bayesian networks underscore the pivotal role of histone modifications in regulating glioma gene expression dynamics.

### Histone Co-Modification Dynamics in pHGGs Validate KDM5B/ARID5B Dysregulation

As shown in Figure 7D, the highest BDM perturbation shifts in the CCLE glioma PID network were observed for the H3K4-H3K56 interaction (30.66 bits), followed by H3K27 and H3K36 acetylation/methylation states. In the BT245 PID network, the top shifts were for H3K4me3-H3K9ac (30.66 bits), followed by H3K9me2-H4K20me3, consistent with the network centrality patterns. For the Spearman network, H3K4me3 and H3K36me3 had the highest eigenvector centrality in BT245, while H3K4me0 and H3K4me1 dominated in CCLE glioma histone networks. These results align with causal metrics like PID and Bayesian inference.

**Figure 7.**
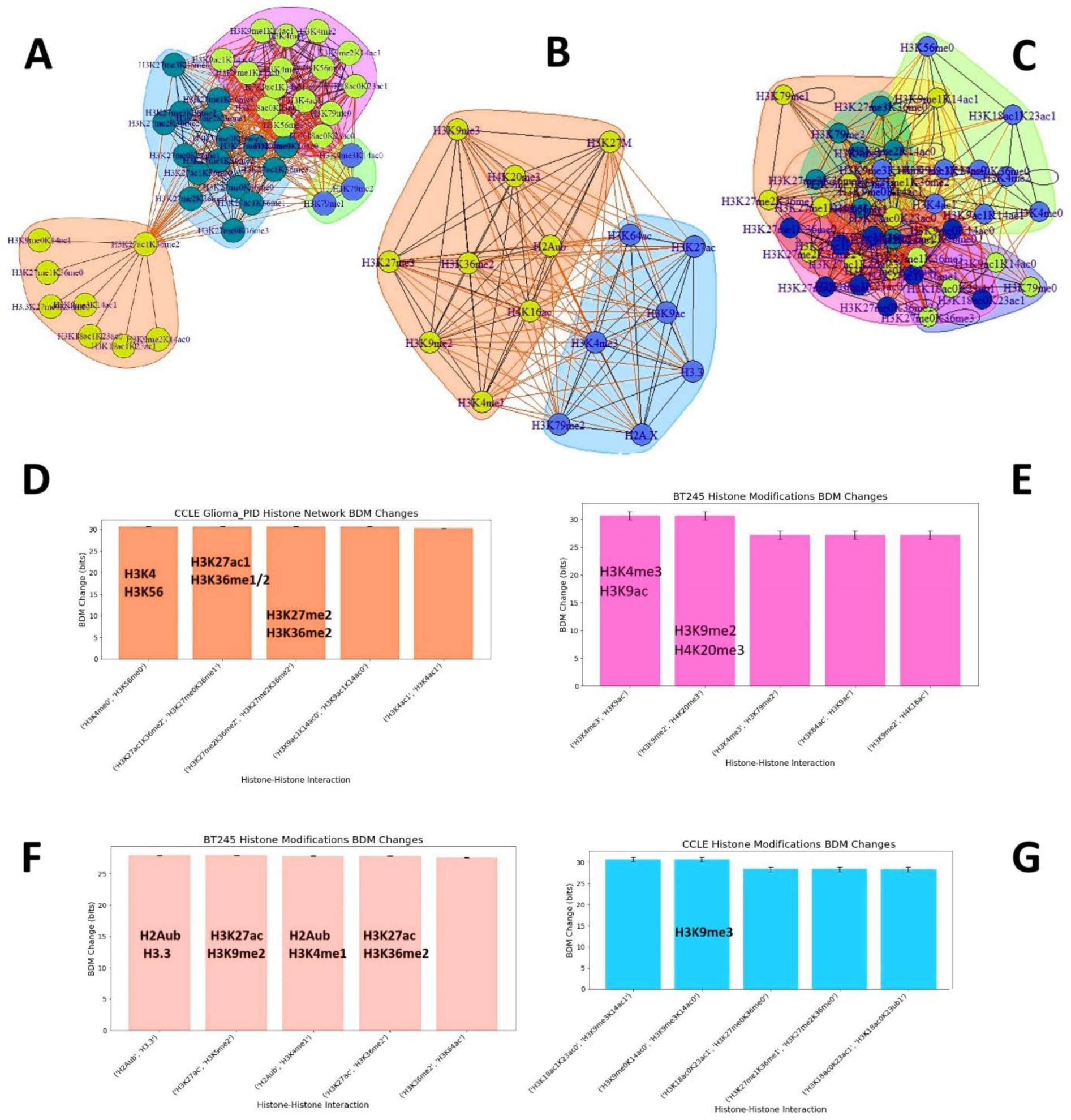
Histone-Histone Modification Networks inferred from CyTOF/ single-cell mass spectrometry reveal H3K4 and H3K9 methylation dynamics as central regulators of pHGG epigenetics. **A)** PID histone co-modification networks from CCLE Glioma datasets. **B)** BT245 histone networks (Harpaz et al., 2022). **C)** Bayesian histone co-modification networks from CCLE datasets. **D-E)** BDM perturbation analysis on CCLE (D) and BT245 (E) PID histone networks. **F-G)** Spearman network centralities for histone networks, highlighting epigenetic plasticity signatures linked to cancer morphogenesis and progression.

BDM perturbation analysis of the BT245 PID network revealed statistically significant interactions between H3K27M and H3K4me3 (Mann-Whitney p=0.032, t-test p=0.024) and H3K27M and H3K9ac (p=0.046 and 0.027). These findings highlight the disrupted repressive epigenetic feedback between H2Aub (ubiquitination via PRC1) and H3K27me3 (via PRC2), a hallmark of K27M gliomas, but also evident in IDHWT glioblastomas without the oncohistone mutations. Fundamentally, K3K4 and H3K9 methylation dynamics, regulated by KDM5B and ARID5B, govern epigenetic and phenotypic plasticity in pHGGs through complex feedback loops and circular causality. These results provide actionable insights for genetic and pharmacological validation of glioma ecosystem regulation.

## DISCUSSION

### Rationale of Complex Systems Modelling

Our methods operate within a *complex systems theory* framework, quantifying emergent behaviors and collective dynamics in single-cell fate decision-making, where the whole system’s interactions reveal more than individual cell fate choices. These methods capture cell fate plasticity dynamics by reconstructing regulatory networks, allowing us to infer cellular trajectories and identify key attractors driving cellular differentiation. Algorithms like scEpath, CellRouter, RBM, and Langevin SDEs utilize probabilistic models, trajectory inference, and attractor dynamics to trace the cellular decision-making process while Bayesian inference and PID-based network analysis reveal the causal relationships controlling these decisions. By modeling these transitions, these methods help decode the underlying molecular mechanisms and potential intervention points in guiding cells toward their favored fate.

### Trajectory Inference Algorithms Predict Neurodevelopmental Patterning Genes and Epigenetic Drivers of Glioma Plasticity

Neurodevelopmental patterning genes such as TGIF1, a regulator of TGFβ signaling, and ZEB1, an EMT driver, were identified across glioma systems, emphasizing their roles in glioma invasion and phenotypic plasticity. HEY1, linked to neuronal-glial fate switches and somitogenesis (William et al., 2007), and PRRX1, critical for mesodermal differentiation (Newton et al., 2022), suggest cross-talk between embryonic somites and neural crest programs. Metabolic reprogramming markers like TSC22D4, HMGB3, and ETV5 reflect adaptive mechanisms of plasticity. These genes highlight the connection between glioma phenotypes and disrupted embryonic development. For instance, the dynamic expression of Hes7 (encoded by the HES7 gene), a marker seen in our CellRouter analysis, is involved in the clock segmentation of somite cells during somitogenesis (Kageyama et al., 2009). Furthermore, FGF and HES7 interactions exhibit oscillatory gene expression dynamics of clock genes/proteins in morphogenesis, which are dysregulated in cancer dynamics (Bessho and Kageyama, 2003; Gibb et al., 2010; Oates et al., 2012; Yoshioka-Kobayashi et al., 2020). Wnt, BMP, EGF, TGFb, and other morphogens observed in our findings act as genetic oscillators coordinating patterning processes in cancer stem cells (Mengel et al., 2010).

KDM5B (H3K4 demethylase) and ARID5B (H3K9me2 demethylase), key epigenetic regulators, were central in IDHWT glioblastoma and are strongly associated with post-resection glioblastoma prognosis, hematopoietic stem cell (HSC) dynamics and leukemia progression, such as acute myeloid leukemia (AML) and lymphoblastic leukemia (ALL) (Boileau et al., 2019; Wang et al., 2020; Leong et al., 2017; McCornack et al., 2023). These epigenetic feedback loops suggest a shared developmental origin for gliomas and pediatric leukemias from the HSC/neural crest bifurcation. H3K9me is involved in transcriptional repression, affecting tumor cell viability, proliferation, and apoptosis, while H3K4me is associated with transcriptional activation, particularly at promoter and enhancer regions, with its modifications playing a key role in GBM stem cell maintenance and differentiation (McCornack et al., 2023). KDM5B influences GATA2 and HOX gene regulatory networks (Cellot et al., 2013), and its dysregulation contributes to high transcriptomic heterogeneity and therapy resistance in glioblastoma (McCornack et al., 2023). ARID5B interacts with the NuRD complex, suggesting it plays a role in transcriptional repression linked to TGFβ and WNT signaling pathways (Kidder et al., 2014). WNT signaling is essential for maintaining GBM stem cell populations, with alterations in histone modifications like H3K4 demethylation, while TGFβ contributes to GBM progression, proneural-mesenchymal transitions (PMT), and immune evasion, with roles in inducing glioma-associated microglia activation and promoting the tumor microenvironment.

Genes like SOX4 (WNT and ERK signaling), HMGB3 (HSC differentiation, WNT signaling), DLX1 (TGFβ regulation), and PRDM9 (H3K4me3 methylation), discovered in this study, are pivotal in glioma plasticity and lineage differentiation. The SCENIC algorithm revealed FOXO1/3 (PI3K/AKT signaling targets) and ARX as enriched regulons in gliomas, highlighting epigenetic vulnerabilities for precision therapies (Wu et al., 2021). Migrasome-associated genes like SEZ6L1 and metabolic regulators such as MEG3 and TSC22D1 were linked to glioma stem cell transitions and plasticity.

### Microtubule Dynamics and Differentiation Markers Regulate Phenotypic Plasticity in pHGGs

As shown in Table 2, the Spearman correlation network analysis revealed **ATF3** as the highest-ranking transcription factor in terms of closeness and eigenvector centrality across both K27M and IDHWT glioblastoma systems, with OTX1 and KLF14 also showing maximum eigenvector centrality (1.00). KLF14, a repressor of TGF-beta signaling, underscores its role in glioma plasticity and embryonic stem cell dynamics. Louvain clustering on KNN graphs identified key plasticity markers: PTPRZ1, PDGFRA, and OLIG1, with significant p-values (e.g., PDGFRA, P = 3.29E-08; OLIG1, P = 3.72E-09), in both glioma systems. Shared immune-inflammatory signatures between the two gliomas included CD74, HLA-DRA, LAPTM5, and RPS4Y1, while IDHWT networks highlighted variations in HLA signatures (e.g., HLA-DQB1, HLA-DRB1) and X-chromosome-linked genes such as XIST and CD99.

**Table 1.** SCENIC Regulons Adjacency Matrix for K27M and IDHWT.

| TF | target | importance | Glioma |
| --- | --- | --- | --- |
| RPL6 | RPL26 | 9.917341 | K27M |
| FOXO1 | MSH4 | 9.719867 | K27M |
| SNAPC4 | ARL2-SNX15 | 8.803067 | K27M |
| ZNF526 | GCSAML | 8.778662 | K27M |
| ZNF99 | PCDHB8 | 8.652782 | K27M |
| MAGED4 | MAGED4B | 8.630525 | K27M |
| FOXD4L4 | FOXD4L2 | 462.6987 | IDHWT |
| MAGED4 | MAGED4B | 459.4231 | IDHWT |
| DLX6 | DLX5 | 124.1313 | IDHWT |
| DLX1 | DLX2 | 117.2777 | IDHWT |

**Table 2.** SCENIC Regulon Networks BDM Shift.

| Gene | Closeness | Eigenvector | Annotations | Network |
| --- | --- | --- | --- | --- |
| ATF3 | 0.043478 | 1 | Proliferation, migration, invasion | IDHWT |
| CEBPD | 0.043478 | 0.988045 | CEPBa; macrophages | IDHWT |
| ATF3 | 0.043478 | 1 |  | K27M |
| BCL3 | 0.043478 | 0.988045 | B-cell leukemia; NF-kB | K27M |
| OTX1 | 0.043478 | 1 | ESC networks | K27M |
| KLF14 | 0.043478 | 1 | Embryonic; TGF-beta receptor II | K27M |

Microtubule-related genes, critical for mitosis and morphogenetic processes, were prominent in both glioma networks. In K27M, TUBB (P = 0.033) and TUBA1B (P = 1.23E-11) emerged as significant markers, while TUBB4A (P = 0.0009) was notable in IDHWT, reflecting its role in neuronal differentiation. The IDHWT glioblastoma network displayed higher expression of neuronal differentiation markers, suggesting a greater inclination toward neuronal lineage commitment compared to the more epigenetically plastic K27M. The top BDM shifts for K27M included interactions such as LAPTM5-MBP (P = 4.01E-06), ERMN-GAS5 (P = 6.68E-07), and CRABP1-EDNRB (P = 7.46E-14), indicating regulatory changes in lineage commitment and plasticity.

Top-ranked BDM changes revealed high-complexity shifts in both glioma systems, with significant interactions such as RPS4Y1-TMSB15A and CNTN1-DKK3 in IDHWT, and GFAP-H2AFZ and PLP1-RGS1 in K27M. As shown in Tables 3 and 4, in K27M gliomas, FABP7, MAD2L1, and TYMS (P = 1.41E-09) emerged as top genes with the most negative BDM shifts, along with interactions involving plasticity genes OLIG1, PDGFRA, and PTPRZ1. FABP7, linked to NOTCH signaling and radial glial fiber formation, is a recognized biomarker in high-grade gliomas. Meanwhile, MAD2L1 underscores the role of mitotic spindle assembly and chromosomal instability. Interestingly, FABP5, associated with metabolic transport and immune signaling, appeared prominently in IDHWT glioblastoma instead of FABP7. Other significant genes included CHL1 (P = 1.69E-06), involved in neuro-cortical development, and DKK3, a WNT pathway tumor suppressor. The divergence between FABP7 in K27M and FABP5 in IDHWT suggests differences in glioma metabolism and lineage plasticity. These findings provide insights into lineage-specific vulnerabilities and therapeutic targets across pediatric gliomas.

**Table 3.** Top Negative BDM changes in CellRouter Spearman Networks.

| Gene1 | Gene2 | BDM Shift | Network |
| --- | --- | --- | --- |
| TYMS | CDK1 | -25.054 | K27M |
| UBE2T | MAD2L1 | -25.265 | K27M |
| OLIG1 | FABP7 | -24.6508 | K27M |
| PDGFRA | FABP7 | -24.4809 | K27M |
| TYMS | MAD2L1 | -24.8499 | K27M |
| MEG3 | PTPRZ1 | -24.755 | K27M |
| LAPTM5 | CD74 | -26.1985 | IDHWT |
| COL21A1 | CFH | -26.4054 | IDHWT |
| DKK3 | CHL1 | -26.4054 | IDHWT |
| MOXD1 | CHL1 | -25.8319 | IDHWT |
| NRCAM | CHL1 | -25.8319 | IDHWT |

**Table 4.** Centrality Measures of CelRouter BDM Perturbation Analysis Networks. DKK3 shows the highest centrality scores in algorithmic complexity shifts for IDHWT glioblastoma.

| Gene | Betweenness | Closeness | Eigenvector | Network |
| --- | --- | --- | --- | --- |
| DKK3 | 31.37687 | 0.017857 | 1 | IDHWT |
| RPS4Y1 | 26.62472 | 0.014085 | 0.830967 | IDHWT |
| CNTN1 | 23.02831 | 0.017544 | 0.968782 | IDHWT |
| PTGDS | 54.83322 | 0.015385 | 1 | K27M |
| MAG | 33.42349 | 0.014706 | 0.957466 | K27M |

**Table 5.** Negative BDM Shifts in RBM and Bayesian PID Networks.

| Network | Dataset | Gene-Gene Interaction | BDM Shift Value |
| --- | --- | --- | --- |
| RBM | K27M | C6orf123, DOCK7 | -30.3 |
| RBM | K27M | KCNJ6, ATG3 | -30.62 |
| RBM | K27M | PTPLB, HHEX | -29.94 |
| RBM | K27M | RLIM, HHEX | -29.94 |
| RBM | IDHWT | DKK3, NMB | -30.42 |
| RBM | IDHWT | TRPM6, GRN | -30.3 |
| RBM | IDHWT | PCDHB12, NOTCH2NL | -29.94 |
| HVG | K27M | ANXA1, IL1B | -32.91 |
| HVG | K27M | APOC1, ADORA3 | -31.3 |
| HVG | K27M | CH25H, AGT | -31.92 |
| HVG | K27M | ADORA3, CXCR4 | -31.92 |
| HVG | IDHWT | RGS1, GAGE8 | -27.06 |
| HVG | IDHWT | TREM2, GAGE8 | -27.06 |
| HVG | IDHWT | C1QA, GAGE8 | -24.97 |
| HVG | IDHWT | GAGE8, CCL3 | -24.76 |
| HVG | IDHWT | GAGE8, CD14 | -24.76 |
| HVG | IDHWT | HLA-DRA, GAGE8 | -24.53 |

**Table 6.** Centrality Measures of Histone PTM Networks. The centrality measures from network adjacency matrices were analyzed using the Mann-Whitney U test, with only the BT245 PID network (Figure 7B, Harpaz et al.) showing statistically significant values (P=0.00018). The CCLE data served as a validation tool, incorporating histone-histone co-modifications to enhance the analysis complex.

| Centrality Measure | CCLE PID Mod. | Value | CCLE Bayes. Mod. | Value | BT245 Mod. | Value |
| --- | --- | --- | --- | --- | --- | --- |
| Closeness | H3K4me1 | 0.02 | H3K9me2k14ac0 | 0.21 | H3K9me3 | 0.18 |
|  | H3K27me0K36me1 | 0.02 | H3K27me2k36me0 | 0.21 | H3K27me3 | 0.18 |
| Betweenness | H3K4me1 | 273 | H3K4me1 | 110 | H3K9me3 | 34 |
|  |  |  |  |  | H3K27me3 | 26 |
| Eigenvector | H3K27ac1k36me2 | 1 | H3K27ac1K36me2 | 1 | H3K4me3 | 1 |
|  |  |  |  |  | H4K16ac | 0.95 |
| Graph Strength | H3K27ac1k36me2 | 142 | H3K27ac1K36me2 | 6.7 | H3K4me3 | 37.86 |
|  |  |  |  |  | H4K16ac | 35.96 |
| Transitivity | 0.7132157 |  | 0.3912402 |  | H4K16ac | 1 |
|  |  |  |  |  | H2Aub | 1 |

### Diverse Stem and Hybrid Cell Fate Lineages in IDHWT Glioblastoma Driven by Neurodevelopmental and Oscillatory Dynamics

CellRouter analysis of IDHWT glioblastoma identified diverse stem cell subpopulations, highlighting a hybrid spectrum of cell states, including mixed OPC/NPC lineages and a unique myeloid-like HSC subpopulation, with key markers such as GATA2, PAX2, and NFE2L2 defining NSCs and TGIF1, IRF8, IFI16, and ATF3 marking lineage plasticity and mesenchymal or immune-inflammatory responses. For instance, NF1-altered glioblastomas demonstrated mesenchymal phenotypes with an upregulation of IRF8, a transcription factor controlling microglial motility, implicating activation of innate immune response pathways (Wang et al., 2021; Larsson et al., 2024). Further, ATF3 is linked to the maintenance of NSC niches and neuronal differentiation programs in the SVZ. (Micheli et al., 2015). Developmental programs like BMP/WNT, TGFβ, and somitogenesis-associated FGF pathways, alongside oscillatory dynamics in genes such as GSX1, HES6, and NFE2L2, control transitions and plasticity within subpopulations. These findings validate ten transcriptional modules previously identified in H3K27M gliomas, highlighting hijacked developmental programs steering glioblastoma stemness, and plasticity transitions.

### Attractor Reconstruction Driven Insights into Glioma Plasticity: Cytoskeletal, Bioelectric, and Neuroimmune Signatures in K27M and IDHWT Subtypes

Gene signatures such as RAP1B, DOCK7, and PAK7 seen in our computational analyses reveal the role of Rho/Rac kinases-based actin dynamics while tubulin genes coordinating microtubule dynamics were repeatedly seen in both pHGG systems. TUBB1A/B is predominant across both K27M and IDHWT systems, as shown by CellRouter networks and scEpath. Gene signatures like CDY2A, SPATA2, and SMARCAL1 are associated with spermatid and testis chromatin remodeling factors, suggesting disrupted developmental pathways in epigenetic regulation. Moreover, DOCK7 reveals that Rho kinases and EGFR signaling may be putative targets for anti-glioma therapies or glioma reprogramming toward stability (non-malignancy). Constitutive EGFR activation promotes glioblastoma invasion through the TAB1-NFKB-EMP1 pathway, while ligand-activated EGFR suppresses invasion via upregulation of BIN3 inhibiting the DOCK7-Rho GTPase pathway, leading to differences in tumor size, invasiveness, and survival (Guo et al., 2022). LAPTM5 is involved in E3, hematopoiesis, immune-inflammatory, and embryonic developmental processes (Glowacka et al., 2012).

The discovery of metabolic signatures such as FOXO3, INPP5K, ATF3, NFE2L2, and CYBB suggest the role of oxidative stress control of mitochondrial bioenergetics in pHGG cellular transitions. We predict that the epigenetic regulators identified by our transcriptomic signatures such as KDM5B, ARID5B, SMARCAL1, PRDM9, FOXA1/2, METTL5, and METTL8, among other chromatin remodeling complexes, steer this epigenetic-bioelectric interdependence in the collective cancer cell fate decision-making.

METTL5 and METTL8 are epitranscriptomic writers involved in catalyzing the m^6^A RNA methylation patterns highly involved in stem cell homeostasis (Boccaletto et al., 2018). For instance, a recent study found METTL1+ neural progenitors as prognostic markers and cell subsets driving IDH-WT glioblastoma (Ji et al., 2023). Many of the BDM signatures and centrality measures such as DKK3 and HHEX, identified WNT as a common target. Further, the emergence of hepatocyte differentiation markers in K27M (including GATA6) speculates that WNT signaling is involved in these cancer developmental processes (Huang et al., 2017). In specific, a hepatocyte-differentiating WNT subtype such as WNT7A, or a subtype involved in embryonic stem cell fate patterning such as WNT5A, may be more specific to reprogramming pediatric gliomas. The emergence of TGIF1, KLF14, ID3, and TGFBI suggests that TGF-b signaling is also a crucial component of glioma cell fate transitions. Signals such as HEY1, CNTN1, FABP7, PSENEN, and NOTCH2NL strongly demonstrate the role of NOTCH signaling in these teleonomic behaviors as well. CNTN1 is also a primary target of PTPRZ1, a highly expressed glioblastoma marker.

To investigate the dynamic structures underlying these patterning processes, we employed RBM, Bayesian networks, and Langevin dynamics modeling as attractor reconstruction methods. RBMs, outperform Hopfield networks for constructing energy landscapes and attractor-based inference, and learn joint probability distributions between visible and hidden (latent) units in a bipartite probabilistic model (Strelioff and Crutchfield, 2014), while Bayesian networks infer outcome probabilities based on dependencies within the network. As shown by the RBM trajectories, critical plasticity genes highlighted cytoskeletal and bioelectric signatures. In K27M, for instance, SPATA2 is involved in microtubule dynamics. Meanwhile, DRAXIN (P=5.27E-13) plays a role in forebrain neocortical neuronal projections, axon guidance, and EMT plasticity. KLF17 and LKLF emerged as common transcription factors to almost 80% of the top importance features extracted by random forests (RF) in g: Profiler gene set enrichment analysis. Interestingly, g: Profiler analysis using only the top 5 importance features revealed WNT, TNF, and SMAD/STAT3 as associated signaling pathways. MTRNR2L1 and MTSS1 (P=6.82E-10) were identified as top-importance features in K27M with RBM.

Meanwhile, the highest RBM importance features in IDHWT were EME2, INPP5K, MEF2BNB, EXOG, DKK3, and DAB2. We also observed SLC3A2 an intracellular calcium and amino acid transporter, and ZNF251 involved in hematopoietic stem cell differentiation. The top 100 importance features predominantly expressed microglia and tumor-associated macrophage (TAM) signatures involved in neuroendocrine regulation and immune-inflammatory processes, including C1QA, CD59, and LAPTM5. The single common importance feature in both systems (K27M and IDHWT) predicted by RBM across the top 100 important features was RLIM (P=0.00014). Multiple algorithms identified RLIM and TERF1 indicating disrupted telomere regulation in the glioma subtypes. RLIM encodes an E3 ubiquitin ligase, a RING-H2 zinc finger protein that plays a role in telomere length-mediated growth controlled by ubiquitination/degradation of TERF1 (Her et al., 2009). TERF1 also emerged as a plasticity signature in glioblastoma.

Endogenous voltage-gated or ligand-gated ion channels such as GRIK3 (kainate receptor subunit), GRIN3B (glutamate ionotropic receptor subunit), NKAIN4, and KCNJ4/6 were also observed among the top 100 RBM-predicted importance features in glioblastoma. Bioelectric networks, through membrane electrophysiology and depolarization, regulate stem cell states and phenotypic plasticity. Levin et al. demonstrated that endogenous bioelectric signals act as "cognitive glue," by coordinating collective cellular decision-making during morphogenesis, with the potential for reprogramming to influence cellular behaviors and therapeutic outcomes (McMillen and Levin, 2024; Kofman and Levin, 2024). TOB1 (a transducer of ERBB2 and BMP), TNFRSF1B (encodes a type of TNF receptor), and CTNND1 were also observed among the importance features. CAMK2D and CDK2 emerged as plasticity signatures, highlighting their role in neurotransmission and neuronal dynamics, as supported by Petralia et al. (2020), who reported upregulated CDK1/CDK2 kinase activity and substrate phosphorylation in pHGG phospho-proteogenomics. These findings highlight the role of the neuro-immune axis and neurotransmission in glioma plasticity.

### Histone PTM Networks Validate KDM5B and ARID5B as Key Drivers of pHGG Cell Fate Decisions

Epigenetic regulation drives GBM phenotypic plasticity through chromatin modifications that shape gene-environment interactions and transcriptional dynamics. In BT245, an H3K27M pHGG cell line, stem-like, fast-proliferating cells show H3K27ac, H4K16ac, and H3K4me1/3, while slow-proliferating, differentiated cells show H3K9me2/3 (heterochromatin and transcriptional repression) (Harpaz et al., 2022). The discovery of ARID5B and KDM5B in glioblastoma suggests that demethylation toward H3K4me1 promotes a more stem-like state due to its role in enhancer activity and cellular plasticity, while demethylation toward H3K9me1 correlates with higher proliferative and less differentiated states. Further, our findings confirm that similar epigenetic drivers are regulating phenotypic plasticity across pHGG systems (i.e., the K27M DIPG and IDHWT glioblastoma). PID network analysis confirmed central histone marks like H3K9me3, H3K27me3, H2Aub, and PRDM9 (H3K4me3/H3K36me3 deposition) as regulators of glioma plasticity. The shared transcriptional circuits and histone modifications (e.g., GATA6 in DIPG vs. GATA2 in IDHWT) suggest a common epigenetic origin for pediatric gliomas, linking their differentiation hierarchies to a broader pediatric cancer network. Harpaz et al. (2022) discovered that the epigenetic heterogeneity within H3K27M gliomas can be characterized by H3K4 and H3K9 methylation profiles.

H3K9me3, H3K27me3, and H4K20me3 are transcriptional inhibition marks linked to X chromosome inactivation and developmental gene signatures like XIST and SATB1, while H3K4me1/3 and H3K27ac activate transcription and regulate differentiation (Liu et al., 2023). These mechanisms dysregulate the expression of developmental gene programs, similar to the disruption of PRC2 altering H3K27me3 deposition in K27M gliomas (Blackledge et al., 2014). ARID5B and KDM5B strongly interact with these histone marks, with KDM5B emerging as a key epigenetic target in leukemias and gliomas (Chan et al., 2022; Ren et al., 2022). Polycomb Repressive Complex 1 (PRC1) dynamics, linked to H2Aub, further highlight this axis in plasticity processes (Zhou et al., 2017; Barbour et al., 2020). In specific, PRC1 leaves the H2AK119ub1 mark of histone H2A at lysine 119. Furthermore, the identification of transition genes and regulons such as GATA6, FOXO3, and MYC in K27M glioma suggests that KDM1A (LSD1) might be involved in their disrupted differentiation processes (i.e., plasticity transitions). Interestingly, KDM1A demethylates H3K4me1/2 and H3K9me1/2 and favors neuronal differentiation programs, suggesting that both investigated pHGG subtypes might be using similar mechanisms to activate a favored lineage transition, or transdifferentiation toward neurons (GeneCard Database; McCornack et al., 2023). This aligns with our previous study, in which using the MuTrans algorithm, we identified transition markers that suggested IDH wild-type glioblastoma exhibited neuronal differentiation programs, microtubule dynamics, and axonal guidance (Uthamacumaran, 2024). This was further supported by g: Profiler gene set enrichment analysis of MuTrans markers (Uthamacumaran, 2024). The Langevin SDE used in this study was inspired by MuTrans (Zhou et al., 2021).

### Convergence of Plasticity Networks: Insights into Glioma Transition Markers and Therapeutic Opportunities

The convergence of distinct algorithmic frameworks towards similar plasticity networks and markers validates the functional interpretations and robustness of our approaches. Furthermore, many previously discovered transition markers (plasticity genes) from our previous study, such as PDGFRA, EGFR targets, OLIG1/2, FXYD6, MTSS1, SEZ6L, MTRN2L1, and SOX11 were also validated in this independent systems medicine analysis. FXYD5 a new variant of the FXYD specific to brain/CNS tissue was also observed. Further, many of the identified transition markers in this study were recently validated as regulatory programs driving phenotypic plasticity in brain tumors using the scRegClust algorithm (Larsson et al., 2024). For instance, Park et al. (2024) identified SOX9, SOX11, and ARID5A among the most critical markers of stemness and proneural-mesenchymal transition (PMT) in glioblastoma. The ERK/MEK and MAPK/RAS pathways identified in our findings suggest the co-existence of diverse developmental programs underlying the lineage-specific plasticity of glioma differentiation. For instance, previous findings show that the EGFR subtype gliomas exhibit a proliferative phenotype with upregulation of key signaling proteins such as CTNNB1 (Wnt signaling), cyclin-dependent genes, and SOX9 (Wang et al., 2021). In contrast, PDGFR subtype gliomas show distinct patterns of PTPN11 phosphorylation with MAPK/RAS and PI3K/AKT downstream signaling. Petralia et al. (2020) found that WNT and GSK3 signaling show upregulation in certain subpopulations of medulloblastoma and low-grade and high-grade glioma.

WNT emerged as one of the central drivers of plasticity networks across both pHGG systems, with signaling from identified transition genes such as HHEX, DKK3, DRAXIN, HMGB3, ARID5B, etc. The WNT subtype is also one of the four major molecular subtypes of pediatric medulloblastoma, the most common brain tumor in children (Northcott et al., 2017; Gold et al., 2024), suggesting there might be overlapping developmental programs driving pHGG subtypes. Further, WNT is critical for leukemogenesis as well (McCubrey et al., 2014). For instance, G34R/V gliomas arise from GSX2/DLX-expressing interneuron progenitors, exhibiting dual astroglial/neuronal identity while actively repressing oligodendroglial programs through impaired differentiation driven by G34R/V mutations and PDGFRA signaling co-option (Chen et al., 2020; Ocasio et al., 2023). We observed GSX1, DLX genes, and PDGFRA as transition markers in both pHGG systems (K27M DIPG and glioblastoma).

Targeting upstream signaling pathways with MEK inhibitors like Trametinib or AKT inhibitors like Ipatasertib may indirectly impact their levels/activity. While previous findings show that the MEK/ERK pathway shows upregulation in a subset of craniopharyngioma tumors and may work better for low-grade gliomas (Petralia et al., 2020), our findings suggest they could be repurposed towards personalized medicine and combination therapies. This also indicates that there might be a feedback loop between MAPK/ERK/MEK pathways and WNT or TGF/Rho signaling regulating plasticity transitions (i..e, from a stem cell to proliferative spectrum). For instance, Studies by Lee et al. (2018) and Gao et al. (2019) demonstrated that chemical cocktails targeting WNT/GSK3β, TGFβ, and Rho Kinase pathways, as well as transcription factors ASCL1 and SOX11, epigenetically reprogrammed adult glioblastoma cell lines into neuronal-like phenotypes (Oh et al., 2017). These neuronal *transdifferentiation* cocktails reduced adult glioblastoma aggressivity by disrupting invasion and proliferation, aligning with the critical phenotypic plasticity genes (transition markers) identified in our data science findings.

### Limitations

Although this study incorporates single-cell data from multiple patients spanning the two pHGG subtypes, the sample size and heterogeneity within and across datasets may limit the generalizability of the findings. Quality control, including data filtering, and preprocessing to address batch effects, noise, and sparsity, was performed using log normalization before analysis. Additionally, computational algorithms like RBM and Bayesian networks rely on assumptions and approximations that may introduce bias or oversimplify complex differentiation processes. Without time-resolved data, potentially from resections in animal models like patient-derived xenografts, porcine glioblastoma models, or organoids, attractor reconstruction can only be inferred. However, the convergence of these frameworks toward consistent plasticity networks and markers lends credibility to the reconstructed patterns. Validation using larger, independent datasets and further integration of multimodal omics data will be critical. Future directions will include advanced deep learning models such as generative AI, graph neural networks, and transformers for pattern discovery in glioma plasticity.

## CONCLUSION

Our multiomics-driven findings suggest that cancers are cellular/developmental *identity disorders* driven by disrupted epigenetic and transcriptional networks. Key mechanisms, including H3K4 and H3K9 demethylase dynamics, immune-microenvironmental and metabolic signatures, microtubule and exosome networks, and telomere-lengthening genes, were found to coordinate phenotypic transitions in both pediatric K27M DIPG and IDH-wildtype glioblastoma, with H3K36 methylation potentially acting as a regulatory switch. Many discovered transition genes, including ATF3, RAP1B, and ARID5B, may play roles in *thanatotranscriptomics* and oxidative stress regulation (Pozhitkov et al., 2017; Antiga et al., 2021). Further, we predict the identified plasticity markers, such as WNT, TGFB, and histone demethylases, are morphogenetic drivers of a developmental program spanning multiple pediatric cancer lineages, including the disrupted neurodevelopmental hierarchies in brain tumors and leukemogenesis, branching from neural crest/tube stem cells and somites (Coste et al., 2015). As suggested by previous studies, our findings also reveal that both pHGG systems consist of a fluid spectrum of stem-cell-like states with hybrid OPC/NPC phenotypes and NSCs as their cell-of-origin (Ocasio et al., 2023; Larsson et al., 2024; Wang et al., 2024; Zhang et al., 2024). Further, the differentiation trajectory from these NSCs seems to activate transcriptional programs favoring a neuronal cell identity due to the retention of stem-cell-like states expressing markers of neuronal lineage.

Our findings reveal that WNT (e.g., WNT7A), Notch (e.g., NOTCH1), and TGFβ morphogen pathways, as seen in Jessa et al. (2022), might regulate dorsal-ventral patterning in the NSCs. In evidence, many of the transition markers identified in our plasticity networks using independent computational frameworks and models are signals associated with WNT pathways. For instance, KDM5B mediated Wnt/β-catenin pathway activation is involved in EMT or PMT, a hallmark of glioma invasion and migration (Yuan et al., 2024). We propose that targeted therapies altering these pathways may help relieve the differentiation blockades. For instance, epigenetic drugs that inhibit KDM5B and ARID5B activity including BAF complex inhibitors (Hodges et al., 2016) or BET inhibitors could be combined with WNT activators and TGFβ inhibitors. Given the complex oscillations driving these patterning systems, the dosing of such combination therapies warrant a more dynamic and adaptive approach. For simplicity, CRISPR-based gene editing or mRNA-based approaches may be more precise reprogramming strategies for preclinical in vitro and in vivo models. Our findings may also pave the way for precision diagnostics and longitudinal monitoring via liquid biopsies for pHGGs through screening migrasome signatures, exosome markers, and plasma-derived, cell-free nucleosome profiling (Harpaz et al., 2022; Fedyuk et al.,2023).

Furthermore, using pharmacological agents to activate or inhibit targeted ion channel types/subtypes involved in pHGG spatiotemporal patterning such as KCNJ4/6, GRIK, and GRIN identified herein, could also allow the phenotypic reprogramming by controlling the bioelectric fields coordinating morphogenesis. Thus, our network signatures serve as potential precision therapies and predictive biomarkers that can reprogram epigenetic marks and restore normal gene expression programs. To conclude, the plasticity networks identified in this study highlight *epigenetic reprogramming* strategies to guide developmentally arrested, maladaptive cancer phenotypes into stable, differentiated states for effective ‘transition therapy’ (Feehley et al., 2023; Yu et al., 2024). Our proposed ‘transdifferentiation therapy’ aligns with psychotherapeutic approaches used in trauma-informed healing, i.e., reintegrating fragmented, disrupted cellular identities stuck from reaching their terminal fate into the *physiological whole-system*, by guiding them toward their natural, desired cell fate commitments.

## Acknowledgments

Dedicated to the children in the B7 Oncology Wing at Montreal Children’s Hospital. I offer my sincere gratitude to Dr. Phil Gold and Dr. Rolando Del Maestro for being *the Light*.

## Declarations

There are no competing interests.

## DATA AND CODE AVAILABILITY

The codes and algorithms for this study can be found in the GitHub repository: https://github.com/Abicu maran/Glioblastoma_III The datasets used in this study can be found in the following papers:

IDHWT glioblastoma scRNA-Seq gene expression counts matrix was obtained from Neftel et al. (2019): https://www.ncbi.nlm.nih.gov/geo/query/acc.cgi?acc=GSE131928

H3.3K27M DIPG scRNA-Seq gene expression counts matrix was obtained from Filbin et al. (2018): https://www.ncbi.nlm.nih.gov/geo/query/acc.cgi?acc=GSE102130

Histone MS Relative Abundance counts for the CCLE glioma samples were obtained from Ghandi et al. (2019):

The CCLE histone dataset can be downloaded from the link below: https://depmap.org/portal/download/api/download?file_name=ccle%2Fccle_2019%2FCCLE_GlobalChro matinProfiling_20181130.csv&bucket=depmap-external-downloads

